# Genomic variants concurrently listed in a somatic and a germline mutation database have implications for disease-variant discovery and genomic privacy

**DOI:** 10.1101/450239

**Authors:** William Meyerson, Mark Gerstein

## Abstract

**Background:** Mutations arise in the human genome in two major settings: the germline and soma. These settings involve different inheritance patterns, chromatin structures, and environmental exposures, all of which might be predicted to differentially affect the distribution of substitutions found in these settings. Nonetheless, recent studies have found that somatic and germline mutation rates are similarly affected by endogenous mutational processes and epigenetic factors.

**Results:** Here, we quantified the number of single nucleotide variants that co-occur between somatic and germline call-sets (cSNVs), compared this quantity with expectations, and explained noted departures. We found that three times as many variants are shared between the soma and germline than is expected by independence. We developed a new, general-purpose statistical framework to explain the observed excess of cSNVs in terms of the varying mutation rates of different kinds substitution types and of genomic regions. Using this metric, we find that more than 90% of this excess can be explained by our observation that the basic substitution types (such as N[C->T]G, C->A, etc.) have correlated mutation rates in the germline and soma. Matched-normal read depth analysis suggests that an appreciable fraction of this excess may also derive from germline contamination of somatic samples.

**Conclusion:** Overall, our results highlight the commonalities in substitution patterns between the germline and soma. The universality of some aspects of human mutation rates offers insight into the potential molecular mechanisms of human mutation. The highlighted similarities between somatic and germline mutation rates also lay the groundwork for future studies that distinguish disease-causing variants from a genomic background informed by both somatic and germline variant data. Moreover, our results also indicate that the depth of matched normal sequencing necessary to ensure genomic privacy of donors of somatic samples may be higher than previously appreciated. Furthermore, the fact that we were able to explain such a high portion of recurrent variants using known determinants of mutation rates is evidence that the genomics community has already discovered the most important predictors of mutation rates for single nucleotide variants.

## Background

Human mutations arise in two major settings: the germline and soma. Germline mutations occur in sperm, eggs, and their progenitor cells and are therefore heritable. Somatic mutations occur in other cell types and cannot be inherited by offspring.

Some somatic and germline mutations matter in health and disease. Critical somatic mutations cause cancer. Somatic mutations have also been known to contribute to autoimmunity [1] and, rarely, seizure disorders [2]. Certain key germline mutations cause heritable disease; and many germline mutations with individually small effects can have a combined [3] impact that becomes meaningful, and which may account for 30-70% [4] of the risk for common diseases.

Nonetheless, in both settings, it is thought that most of the variants that arise are neutral [5] or nearly neutral [6], the result of stochastic mutational processes that alter the genome. In the germline, which is relatively shielded from the environment, the most active mutational processes are endogenous to the cell, such as errors in DNA replication and spontaneous DNA damage [7]. In the somatic setting, additional environmental exposures, such as ultraviolet light [8] and cigarette smoke are active. Cancerous somatic cells are frequently further deranged with cancer-specific defects in DNA repair as a further source of mutations [9]. While some of these mutational processes can be identified, others can only be inferred on the basis of mutational signatures – patterns in the spectrum of mutations present across samples [10]. Mutational processes differ in prevalence across samples, and within samples they have a predilection for certain nucleotide contexts. Altogether, genomic sites differ in mutation rate depending on the features of those sites, the internal state of the cell [11], and environmental exposures.

The main challenge in identifying and interpreting somatic and germline variants with the largest impact in disease is the sheer number of neutral and nearly neutral variants from which they must be distinguished [12]. Thus, one core priority in biomedical genomics is to understand the patterns these neutral variants take and the mutational processes that lead to them. This goal complements activities in evolutionary genomics, which seeks to understand how mutation and natural selection have led to the diversity of life.

In order to better understand the patterns and processes of mutation, there are two kinds of facts about the relationship between somatic and germline variants that are useful to learn, for reasons that will become clear. First, it is useful to know *how* similar somatic and germline mutation patterns are to each other. Second, it is useful to know *why* these patterns have the degree of similarity that they do.

A key reason that it is useful to know how similar somatic and germline mutation patterns are to each other has to do with building better models of background mutation rates. Models of background mutation rates are used to distinguish disease-causing mutations that are subject to evolutionary selection pressure from the range of neutral and nearly neutral variants that occur by chance. The accuracy of these models depends on the number of variants used. One way to acquire more variants for these models is to aggregate variants across patient samples. However, it is only meaningful to aggregate variants across samples that have similar mutation patterns. The largest databases of human variants in existence are separated into somatic and germline call-sets. If somatic and germline mutation patterns are sufficiently similar, it may be possible to aggregate somatic and germline variants together to build very precise background mutation models against which to better distinguish disease-causing mutations. Building these combined models is a formidable task beyond the scope of this paper, but the first step is to lay the groundwork for this possibility by assessing the total similarity between somatic and germline mutational processes.

Understanding why somatic and germline mutation patterns agree with each other to the extent that they do is useful for many reasons. First, and most simply, this could highlight mutational processes that are shared between the soma and germline. Second, the combined model-building described above should be guided by a deeper understanding of the relationship between these mutation patterns so that variants can be aggregated in a principled way that respects the similarities and differences of somatic and germline mutation patterns. Third, attempting to understand why somatic and germline patterns agree can reveal how much similarity *cannot* be accounted for by known factors. This is useful to know because it suggests how much or what kinds of unknown determinants of mutation rates are out there waiting to be discovered. Fourth, one explanation for similar patterns in somatic and germline call-sets is of special importance: germline leakage, the misclassification of germline variants as somatic. Germline leakage is a concern both for technical applications and also for the genomic privacy of somatic donors [13].

A number of studies have been conducted comparing somatic and germline variant patterns which have bearing on some of these goals. Milholland *et al*. found that the mutation rates of the basic types of nucleotide substitutions are moderately correlated between the soma and germline [14]. Hodgkinson *et al.* compared the somatic mutation rates by megabase from 3 cancer patients with human-chimp divergences and found modest correlations by genomic region in somatic and germline mutation rates [15]. Chen *et al*. showed that epigenetic features associated with higher (or lower) mutation rates in the soma tend to be associated with higher (or lower) germline mutation rates [16]. Rahbari *et al*. investigated mutational signatures and argued that somatic Signatures 1 (spontaneous deamination of 5-methylcytosine) and 5 (putative but unknown endogenous mutational process) are the only somatically-derived signatures detectable in both the germline and soma [17].

One approach that has been missing from the literature is a focus on single nucleotide variants that are concurrently mutated in the soma and germline. Recurrence of single nucleotide variants is attractive because it implicitly aggregates the effect of many of these individual determinants found by other studies, can include the effects of undiscovered determinants [18], and provides a convenient way of testing the impact of the different determinants of mutation rates. It is also particularly well-suited to studying germline leakage because that occurs at a single nucleotide scale.

A historic barrier to single nucleotide variant (SNV)-level recurrence approaches is their requirement for a high density of variants called with technical uniformity. This barrier has recently been lifted by new data sets. Our SNV-level analysis is made possible by two vast, uniformly re-called public data sets of somatic and germline variation: the genome Aggregation Database (gnomAD) [19] of 120,000 germline whole exomes and The Cancer Genome Atlas (TCGA) [20] of 10,000 somatic whole exomes from cancer patients, which were recently harmonized across cancer types [21].

## Results

### Simulated impact of shared mutational processes on shared SNVs

Before calculating the number of SNVs concurrently mutated between the soma and germline (cSNVs), we wanted to establish a baseline for interpreting these numbers. Therefore, we conducted simulations to test how unknown mutation processes of various kinds would affect the number of shared SNVs (Methods). A simulated mutational process increases or decreases the mutation rate of the same random subset of genomic sites in the soma and germline. Excess recurrence was detectable either when the fraction of affected bases was moderately high or when the mutational process had a large effect on mutation rates (Figure 1). From these simulations, we calculated that a mutation process that affects about 1% of the genome must increase the mutation rate of the affected bases about 4-fold to increase the number of recurrent variants by 5% above expectations. Our power analysis (Methods) indicates that with current sample sizes, we are theoretically powered to detect a 0.6% excess of recurrent variants; once somatic databases grow to the size of current germline databases, a 0.35% excess recurrence could be detected. In contrast, if we were to rely on publicly available *de novo* variants, the minimal detectable excess of cSNVs would need to be 12% above expectations, due to their smaller sample size. Both mutation-promoting processes and mutation-inhibiting processes led to excess recurrence. These simulations show that recurrence analysis is theoretically well-powered to detect the impact of a broad range of shared mutational processes that might be active in our somatic and germline data sets.

**Fig. 1:**
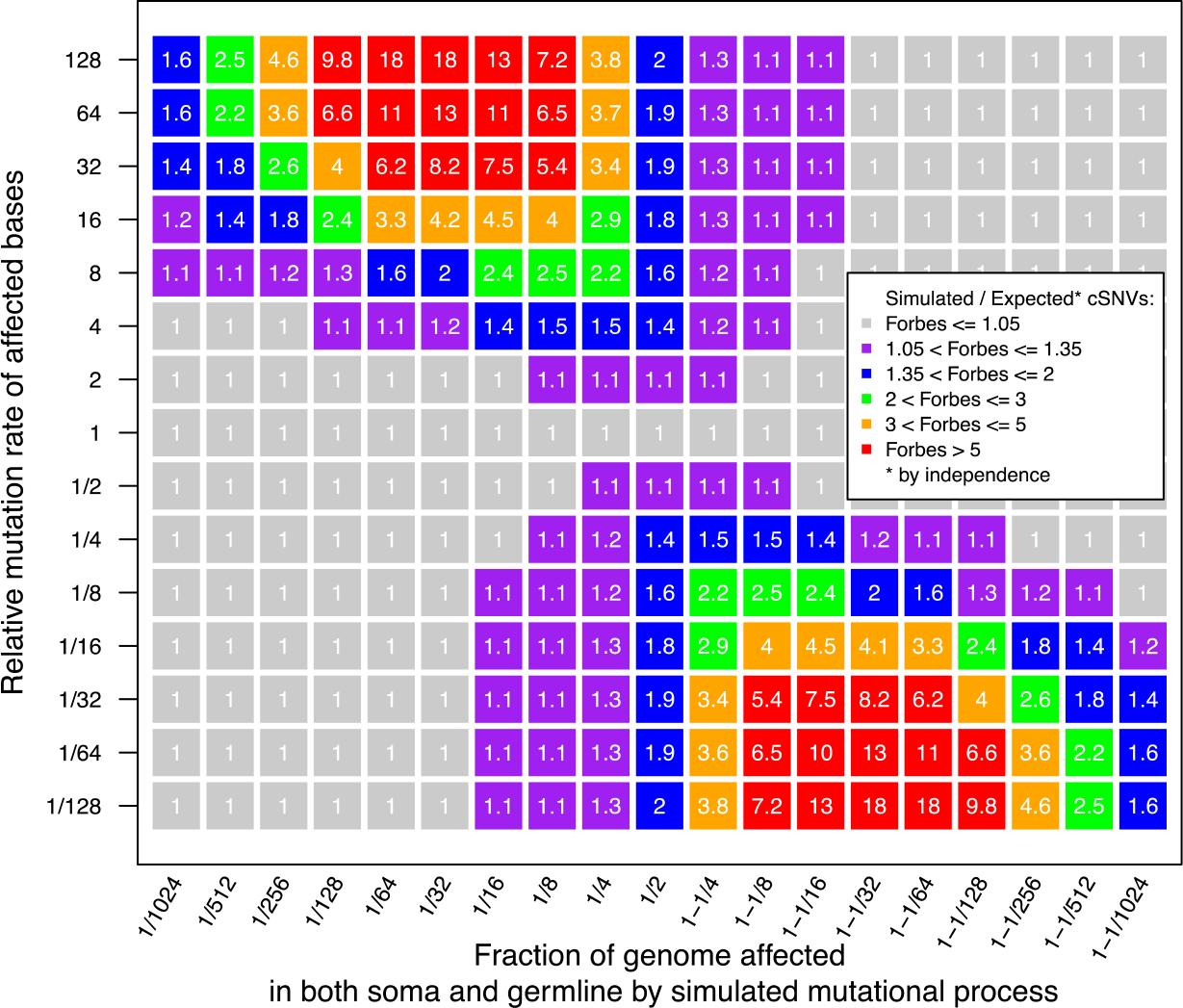
Simulated mutational processes generate a detectable excess of recurrent variants Unknown, hypothetical mutational processes were simulated to act in a coordinated manner in the soma and germline. Each simulated mutational process equally affects the same effectively random subset of genomic sites in the soma and germline. This leads to an excess of sites that are concurrently mutated in the soma and germline, over expectations by independence (the Forbes coefficient). The magnitude of resulting Forbes coefficients (numbered grid squares) depends on the fraction of genomic sites subject to the mutational process (x-axis) and the mutation rate multiplier (y-axis) of the affected bases relative to unaffected bases. Symmetry arises because a mutation-promoting process affecting 25% of the genome is equivalent to a mutation-inhibiting process affecting 75% of the genome.

### Many more variants shared between the soma and germline than expected

Our first goal was to calculate the enrichment of cSNVs over expectations (the cSNV Forbes coefficient [22] – see Methods). This quantity has a technical and biological component. To enrich for the biological component, we filtered genomic sites to create a conservative whitelist of the most technically unproblematic sites. These sites exclude genomic regions known to be prone to technical errors and are uniquely mappable, well-covered, GC-normal, and free of common germline polymorphisms. 16,879,845 genomic sites pass all filters, implying a universe of 50,639,535 potential SNVs. Of these, 3,339,715 are observed in the germline database (gnomAD) and 1,309,369 are observed in the somatic database (TCGA). Under statistical independence, it was expected that 86,354 unique SNVs would be simultaneously present in gnomAD and TCGA; instead, we observed 268,250 concurrent variants, a 3-fold enrichment (Forbes coefficient 3.106, binomial p-value << 5e-324) in our maximally-filtered set. This overall enrichment was not sensitive to filtering strategy: with minimal filtering, this coefficient is 2.95. The calculated cSNV Forbes coefficient on the filtered set represents our first-pass estimate of the total similarity between somatic and germline mutation patterns. Our simulations indicate that there are many ways this enrichment could arise (green and orange grid elements of Figure 1); one example is if the same quarter of the genome was 16-fold more mutable than the rest of the genome in both the germline and soma.

To assess the representativeness of these findings, we calculated the cSNV rate separately for each somatic sample, where the cSNV *rate* of a set of variants from one database is the fraction of those variants that co-occur in the second database. We observed that an elevated cSNV rate is a pervasive phenomenon, not confined to a few outlier somatic samples (Figure 2).

**Fig 2.**
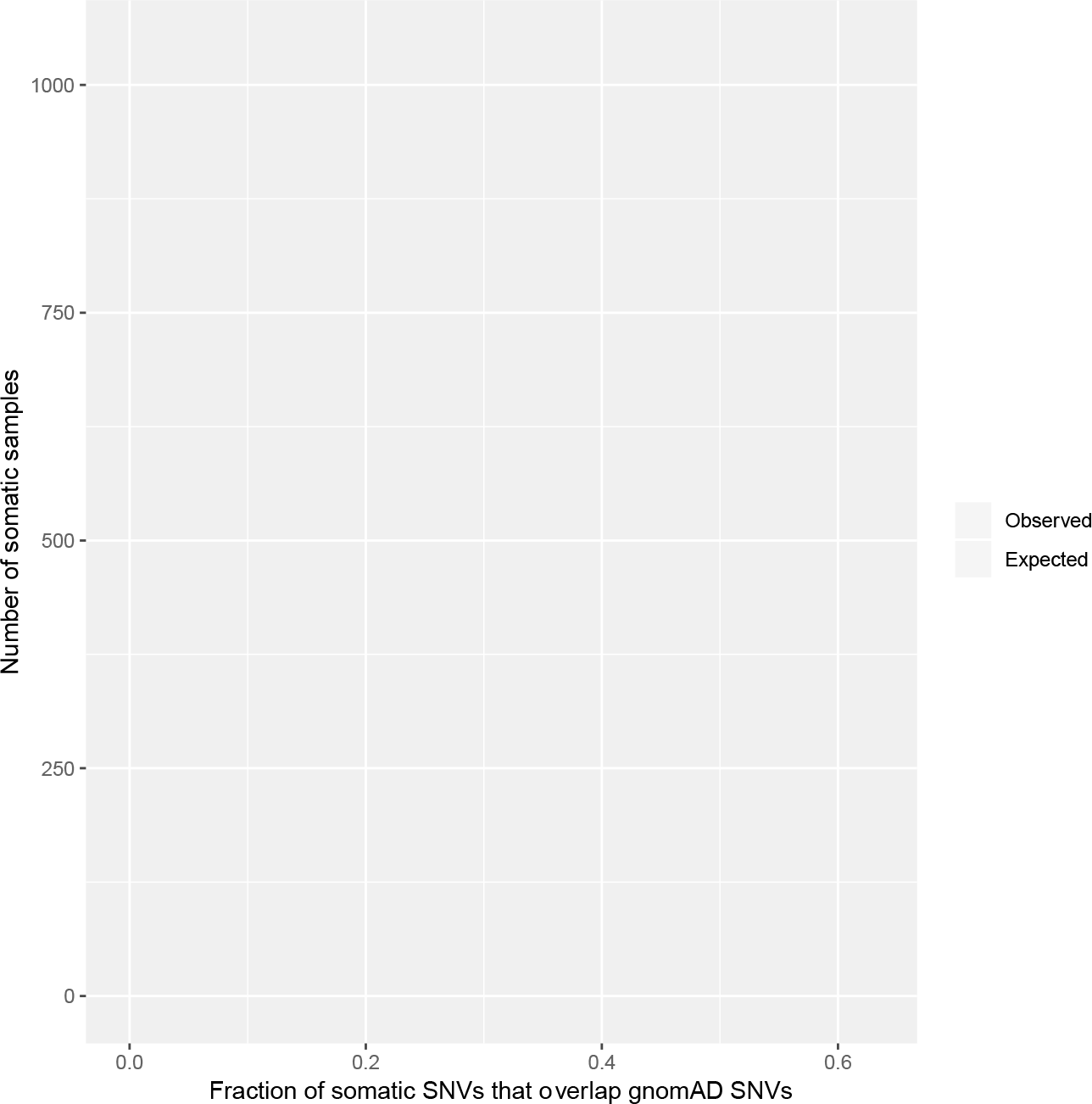
Excess cSNVs are broadly distributed across samples These histograms show the observed (purple) and expected (pink) distributions of cSNV rates across somatic samples. The observed distribution is strongly right-shifted compared with expectations. Here, expectations for each tumor are modeled using a binomial draw, setting the number of trials equal to the number of somatic SNVs in the tumor and the success probability equal to the constant average germline mutation rate. The peak at 0 in the expected distribution reflects the fact that enough tumors have a very small number of mutations that it would be expected that some of these tumors would have 0 cSNVs.

### cSNV enrichment reproduced in non-cancerous somatic samples

We also repeat the analysis using a set of 385 white-listed SNVs taken from normal somatic stem cells [23]. The Forbes coefficient between gnomAD and these normal somatic SNVs was 4.53, indicating that the overlap between somatic and germline samples is not unique to cancer, and may be enhanced in normal tissues, which could reflect the lower burden of cancer-specific mutational processes in normal tissues.

### Quantity and quality of insertion and deletion calls insufficient for recurrence analysis

11,909 somatic insertions and 2,417 rare germline insertions pass our filters. It was expected that 1.7 of these would have the same start coordinate; instead 27 were observed. Of these, 22 are flanked by homopolymer repeats of length 4 or greater, which suggests involvement of strand slippage either of the *in vivo* DNA polymerase or in the sequencing reaction.

36,210 somatic deletions and 6,314 rare germline deletions pass our filters. It was expected that 13.5 would have the same start coordinate; 9 were observed. Because of the low counts of expected and observed insertions and deletions concurrently mutated in the soma and germline, we focused the remainder of the analysis on SNVs.

### Some cancer types share variants with the germline more frequently than do others

The cSNV enrichment was also calculated separately for each cancer type. The observed number of cSNVs exceeded expectations in all cancer types, with a minimum Forbes coefficient of 1.62 in lung adenocarcinoma and a maximum coefficient of 5.81 in uveal melanoma (Supplemental Table 1)(all binomial p-values < 2e-18). The Multi-Center Mutation Calling in Multiple Cancers (MC3) project, completed in March 2018, harmonized the TCGA cancer cohorts with each other to minimize technical differences between them [21]. The different cancer types are, however, subject to different biological sources of mutation; thus, the large variation in cSNV enrichment by cancer type suggests that biological factors play some role in cSNV enrichment.

### Overview of the explanatory model

Our next goal was to explain the observed excess of cSNVs. Existing binary similarity measures, such as the Forbes coefficient of association *F* = *c/e* (where *c* is the number of observed cSNVs and *e* the number of cSNVs expected by independence), are well-suited for quantifying departures of observed cSNV counts from expectations under statistical independence. However, to explain departures, we developed a statistical framework (Methods) that estimates the portion of excess cSNVs that may be accounted for by categorical variables associated with both somatic and germline mutation rates.

Our measure, the partitional dependence of *F* on categorical variable *v* (such as nucleotide context), is given by

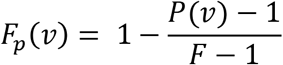

where *P(v)*, the partition-conditioned Forbes coefficient is given by

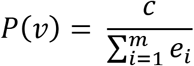

where *c* is the number of cSNVs, *m* is the number of factor levels of categorical variable *v* and

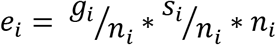

and where *g_u_ s_i_*, and *n_i_* are the number of G events, S events, and elements of the *i*th partition of the full domain.

When *F_p_*(*ν*) = 0, variable *v* does not account for any observed excess or shortage of shared variants. When *F_p_(ν)* = 1, variable *v* can account for all of the observed excess or shortage of shared variants.

### Nucleotide context is a major determinant of variants shared between the soma and germline

We tested how well nucleotide context can explain excess cSNVs. We initially observed that 67% of all cSNVs occur at N[C->T]G contexts, which are known [24] to frequently mutate in the germline and soma. Our Forbes dependence metric estimated that 92% of the cSNV enrichment may be attributed to the high rate of N[C->T]G mutations. Our local Forbes coefficient test estimated that N[C->T]G cSNVs and non- N[C->T]G cSNVs occur 4% and 55% more frequently than expected, respectively, after conditioning on the base rates of these particular kinds of mutation in the germline and soma. Further partitioning potential SNVs into seven types of context-related variants (see Table 1 legend) explains 97.2% of cSNV enrichment. Extended nucleotide contexts added minimal explanatory value overall, but offered a moderate boost in the ability to explain cSNVs outside of N[C->T]G contexts (from 82.5% using seven types of nucleotide contexts to 88.6% by treating each of 24,576 heptamers separately).

**Table 1:** Excess recurrence is statistically explainable by nucleotide context

| Nucleotide context | Number of partitions | Partition-conditioned score | Partition dependence | Partition-conditioned score, excluding N[C->T]G | Partition dependence, excluding N[C->T]G |
| --- | --- | --- | --- | --- | --- |
| Unpartitioned | 1 | 3.106 | NA | NA | NA |
| N[C->T]G vs all others | 2 | 1.168 | 92.0% | 1.553 | NA |
| Seven Type | 7 | 1.059 | 97.2% | 1.097 | 82.5% |
| Trinucleotide | 96 | 1.064 | 97.0% | 1.098 | 82.3% |
| Pentanucleotide | 1536 | 1.054 | 97.4% | 1.071 | 87.1% |
| Heptanucleotide | 24,576 | 1.051 | 97.6% | 1.063 | 88.6% |

Seven Type refers to the 7 basic types of variant from Arndt *et al*. (C->A, C->G, C->T, N[C->T]G, T->A, T->C, and T->G after collapsing purine-centered contexts onto their central pyrimidine) [40]. The partition-conditioned score gives the excess recurrence over a form of expectations that incorporate the fact that different nucleotide contexts have different mutation rates. The partition dependence gives the percent of excess recurrent variants that can be statistically explained by the different mutation rates of the various partitions. See Methods for details.

Some pentamers were found to carry more cSNVs than expected after taking into account the mutation rate of these pentamers in the soma and germline. The most well-powered outliers are listed in Table 2, below. Three out of four of these outlier pentamers contain a C->A mutation within a T-rich context.

**Table 2:** Outlier pentamers with high recurrence rates not explained by their average mutation rates

| Pentamer | Germline cnt | Somatic cnt | Reference cnt | cSNV cnt | Local Forbes Coef |
| --- | --- | --- | --- | --- | --- |
| AT[C->A]TT | 2236 | 5229 | 45328 | 374 | 1.45 |
| CA[T->C]TG | 18256 | 1337 | 42592 | 752 | 1.31 |
| TT[C->A]TC | 1734 | 8067 | 60250 | 385 | 1.66 |
| TT[C->A]TT | 2914 | 11887 | 68309 | 821 | 1.62 |

The success of nucleotide context at explaining cSNV rates implies that nucleotide contexts have correlated mutation rates in the germline and soma, as observed in [14]. We next tested this explicitly. The somatic and germline mutation rates by nucleotide context were substantially correlated (Table 3). They remained correlated at extended nucleotide contexts, which is a novel finding, even after controlling for the central bases.

**Table 3:** Correlations between somatic and germline mutation rates by nucleotide context

| Nucleotide Context | Somatic-Germline Spearman correlation coefficient | P-Value |
| --- | --- | --- |
| Trimer | 0.85 | < 2.2e-16 |
| Trimer, excluding NCG sites | 0.83 | < 2.2e-16 |
| Pentamer | 0.84 | < 2.2e-16 |
| Pentamer, controlling for central 3 bases | 0.31 | < 2.2e-16 |
| Heptamer | 0.74 | < 2.2e-16 |
| Heptamer, controlling for central 5 bases | 0.15 | < 2.2e-16 |

### Signature analysis

Signature 1 is a ubiquitous pattern in the mutational spectra across cancers; its presence is taken to imply the existence of an underlying mutational process that introduces mutations with the spectra seen in Signature 1. The mutational process that is the source of Signature 1 has been previously identified as the spontaneous deamination of 5-methylcytosine [25]; this deamination is known to be active in the germline as well. We show that the activity of Signature 1 in both the soma and germline helps to explain excess cSNVs.

We tested whether the prevalence of the various mutational signatures per somatic sample could predict the cSNV rate by somatic sample. These prevalences were obtained from Huang *et al*. [26] The most significant correlation was that the prevalence of Signature 1 by somatic sample predicts 31% of the variation of cSNV rates by somatic sample (p value < 2e-16) (Figure 3), which reinforces recent findings that the endogenous mutation process Signature 1 is active in both the soma and germline. Similarly, the median prevalence of Signature 1 by cancer type was strongly correlated with the median cSNV rate by cancer type (Figure 4). In this analysis, cancer types at the extremes can be rationalized: perhaps lung adenocarcinoma (LUAD) is less germline-like because cigarette smoke introduces a very different spectrum of mutations than do endogenous processes, and perhaps glioblastoma multiforme (GBM) tumors are more germline-like because they are encased within the skull and therefore mostly subject to endogenous mutational processes.

**Fig 3.**
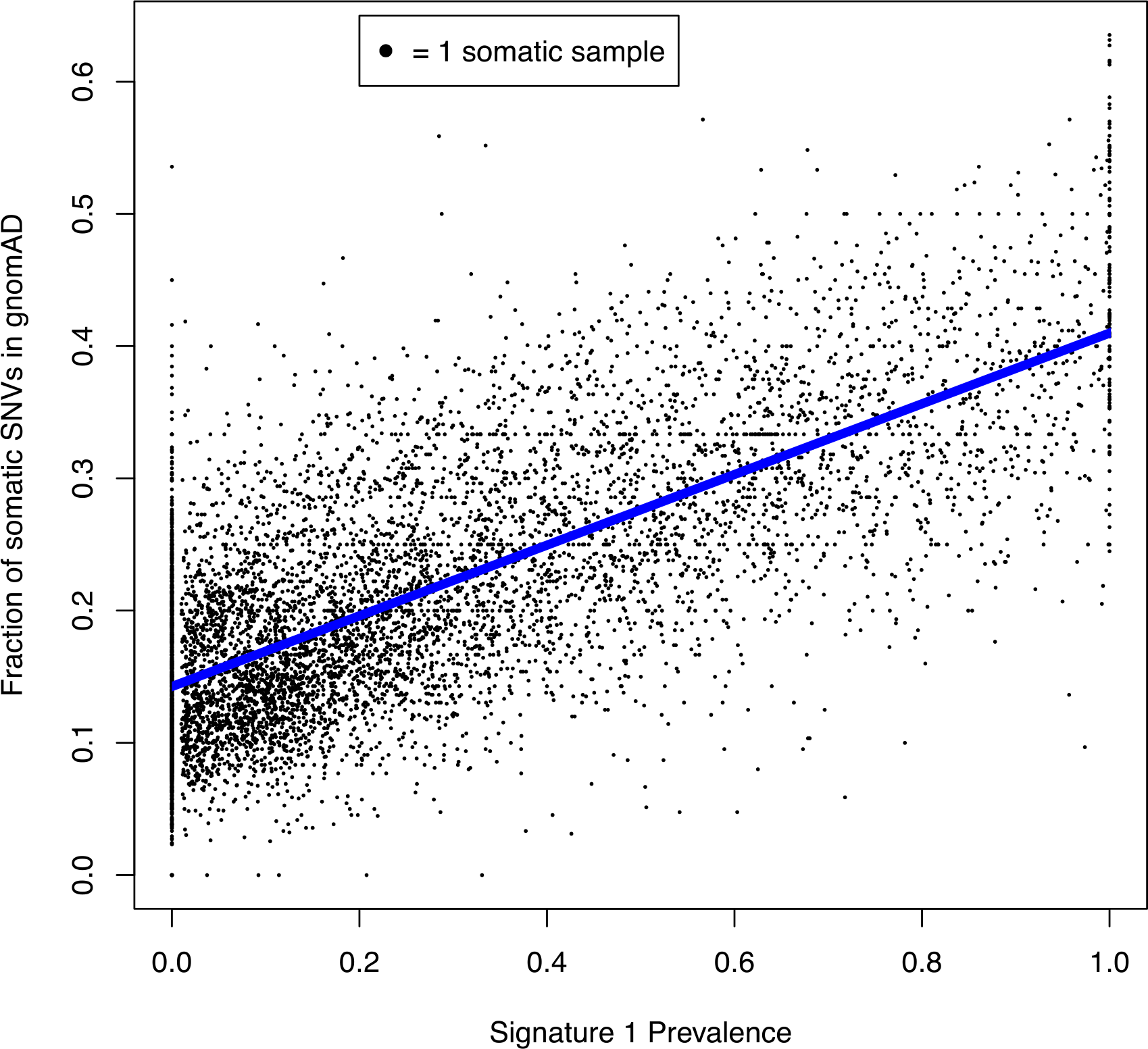
The prevalence of Signature 1 in a somatic sample predicts its cSNV rate Each point of this scatterplot represents a somatic sample. On the x-axis is the proportion of SNVs in that sample that are attributed to mutation Signature 1. On the y-axis is the proportion of SNVs in that sample that can also be found in the germline database. Somatic samples with fewer than 20 total SNVs are excluded. The trend line is shown in blue: somatic samples with a higher prevalence of Signature 1 have a higher rate of cSNVs.

**Fig 4.**
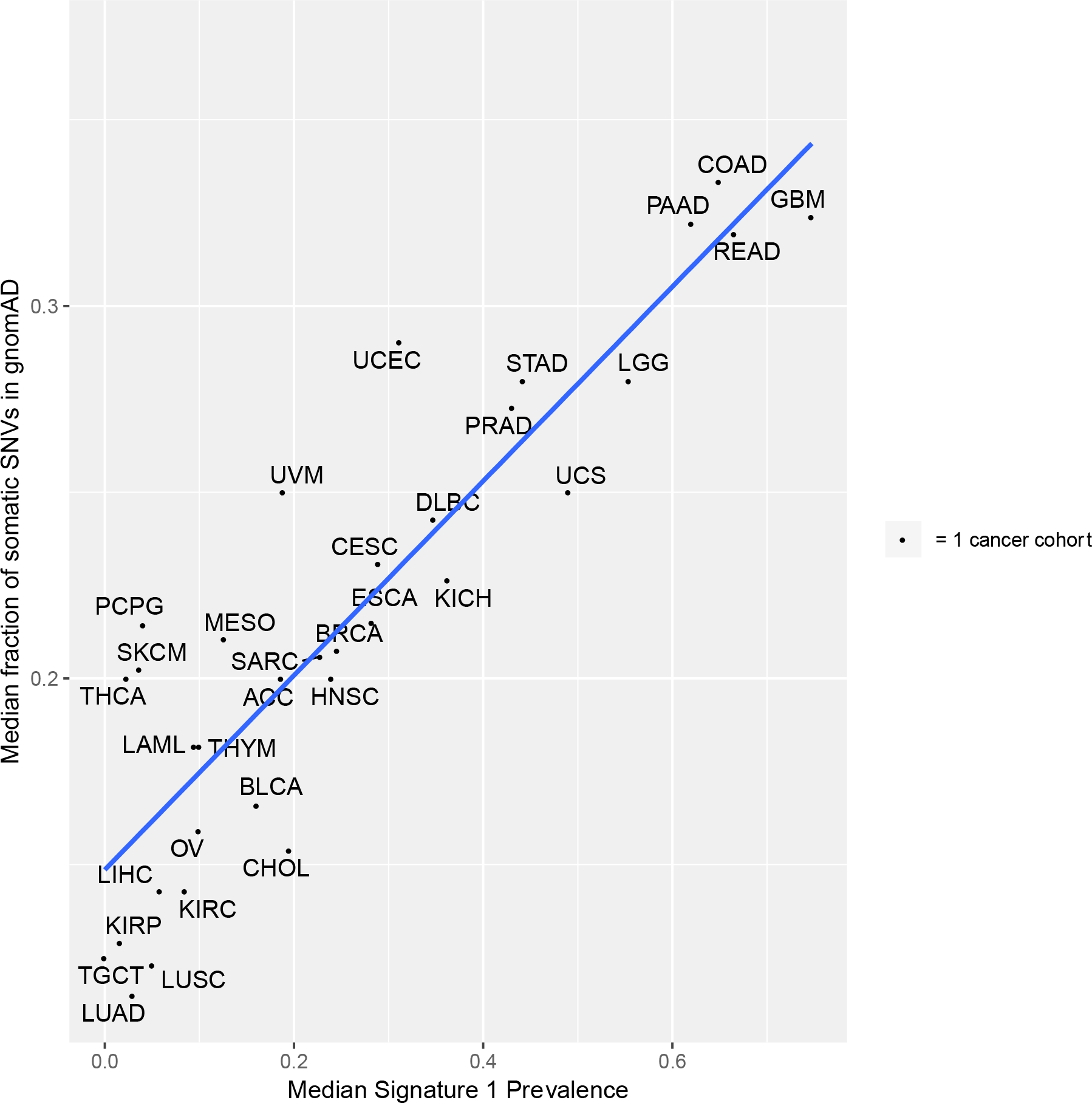
The prevalence of Signature 1 in a TCGA cohort predicts its cSNV rate Each point represents a cancer cohort. On the x-axis is the proportion of SNVs in that sample that are attributed to mutation Signature 1. On the y-axis is the proportion of SNVs in that sample that can also be found in the germline database. Cancer types are labeled according to the official TCGA codes [41].

Signature 5 is a common pattern in the mutational spectra across cancers. No underlying mutational process has been conclusively identified as the source of Signature 5; therefore, it is not conclusively known whether that process is also active in the germline. In [17], it was argued that, based on the mutation spectra in the human germline, Signature 5 appears to be active in the human germline as well. Somewhat unexpectedly, we find that, if Signature 5 is active in both the soma and germline, it does not help explain excess cSNVs. Specifically, we found that the prevalence of Signature 5 across somatic samples was not positively associated with the samples’ cSNV rates. There are many potential reasons why it might not: Either Signature 5 is not as active in the germline as Rahbari *et al*. estimated, or its action is diffuse across many genomic sites, or its prevalence in somatic samples comes at the cost of Signature 1, which is even more tightly linked to germline mutational processes than is Signature 5.

The only other signatures whose prevalences were found to significantly positively associate with cSNV rates were Signatures 6 and 15 with R-squared 0.024 and 0.012, respectively (p value < 2.2e-16). There is no known association of Signatures 6 or 15 with the human germline; however, Signature 6 and Signature 15 are the signatures whose nucleotide spectrum is most similar to Signature 1 (Pearson’s rho 0.81 and 0.48, respectively). These results suggest that mutation rates by nucleotide context are correlated between the soma and germline largely because of their shared exposure to Signature 1, with a minor component from similarities certain somatically-exclusive signatures have with Signature 1.

### Genomic region is a minor determinant of variants shared between the soma and germline

The effects of genomic region were minor. Similarities in the somatic and germline mutation rate by megabase explains only 0.4% of excess cSNVs. (Table 4)

**Table 4:** Genomic region explains a small fraction of excess cSNVs.

| Region | Number of partitions | Partition-conditioned score | Partition dependence |
| --- | --- | --- | --- |
| Whole Genome | 1 | 3.106 | NA |
| Chromosome | 22 | 3.107 | 0.0% |
| Megabase | 2394 | 3.097 | 0.4% |
| 100kb | 14,214 | 3.084 | 1.0% |
| 10kb | 54,889 | 3.076 | 1.4% |

Nonetheless, combining regional features with nucleotide context features explained a greater share of excess cSNVs (97.9%) than did nucleotide context alone (97.2%) (Table 5). The increased explanatory power of the combined model was not an artifact of the greater number of partitions in the combined model, because a dummy combined model, which randomized the megabase membership of SNVs, did not lead to any change in explanatory power compared to the nucleotide context-only model (97.2% in both cases).

**Table 5:** A combined model with region and nucleotide context explains excess cSNVs slightly better than nucleotide context-only model

| Bin | Number of partitions | Partition-conditioned score | Partition dependence |
| --- | --- | --- | --- |
| Whole Genome | 1 | 3.106 | NA |
| Seven Type | 7 | 1.059 | 97.2% |
| Megabase | 2394 | 3.097 | 0.4% |
| 100kb | 14,214 | 3.084 | 1.0% |
| 1MB x Seven Type | 16,737 | 1.054 | 97.4% |
| 100kb x Seven Type | 98,960 | 1.044 | 97.9% |
| Dummy.100kb x Seven Type | 99,250 | 1.058 | 97.2% |

### Germline contamination appears to be active but predictively redundant

We used two independent strategies for assessing the extent of germline contamination among cSNVs. First, we re-computed the cSNV enrichment rate after replacing gnomAD with denovo-db [27] as the germline database with which to intersect with TCGA. Denovo-db (version 1.6) comprises 7,296 unique SNVs that have been confirmed through parental-offspring trio sequencing to have arisen independently in the germline donor and pass our stringent filters. *De novo* germline variants are free of the past inheritance structure that makes accumulated germline variants prone to contaminating somatic samples. We find that the cSNV enrichment rate is remarkably higher using this database of *de novo* variants (Forbes coefficient 5.11), indicating that germline contamination does not explain a large fraction of the cSNVs observed using gnomAD.

It is unclear why the Forbes coefficient is so much higher when using *de novo* variants, but a similar enrichment is seen among germline variants present in 5-40 unique gnomAD individuals, consistent with a depressed gnomAD Forbes coefficient being a consequence of binarization of gnomAD allele counts [28] (see Methods).

Second, we used a read depth analysis to quantify the amount of germline contamination and its impact on the cSNV rate. TCGA makes use of matched normal samples to reduce germline contamination when calling somatic variants; that is, for a variant to be called in TCGA as a somatic variant, it must not be detected in the germline of the same patient after sampling at least 8 reads from the normal sample for that patient at that site. Because of the stochasticity of sequencing, the ability of this filter to remove germline contaminants depends on the read depth in the matched normal sampled at the site that the somatic variant is called. We find that increased read depth in the matched normal is associated with a decreased cSNV rate in gnomAD, from 29.4% at a read depth of 8, leveling off to a gnomAD cSNV rate of 21%-22% at a read depth of 100 and above. (Figure 5) The leveling off of the cSNV rate at increased normal depth suggests that there is no appreciable germline contamination when matched-normal read depth exceeds 100. The fact that there is a higher cSNV rate at lower matched-normal read depths suggests that germline contamination inflates the cSNV rate for TCGA variants with low depth in the normal.

**Fig. 5:**
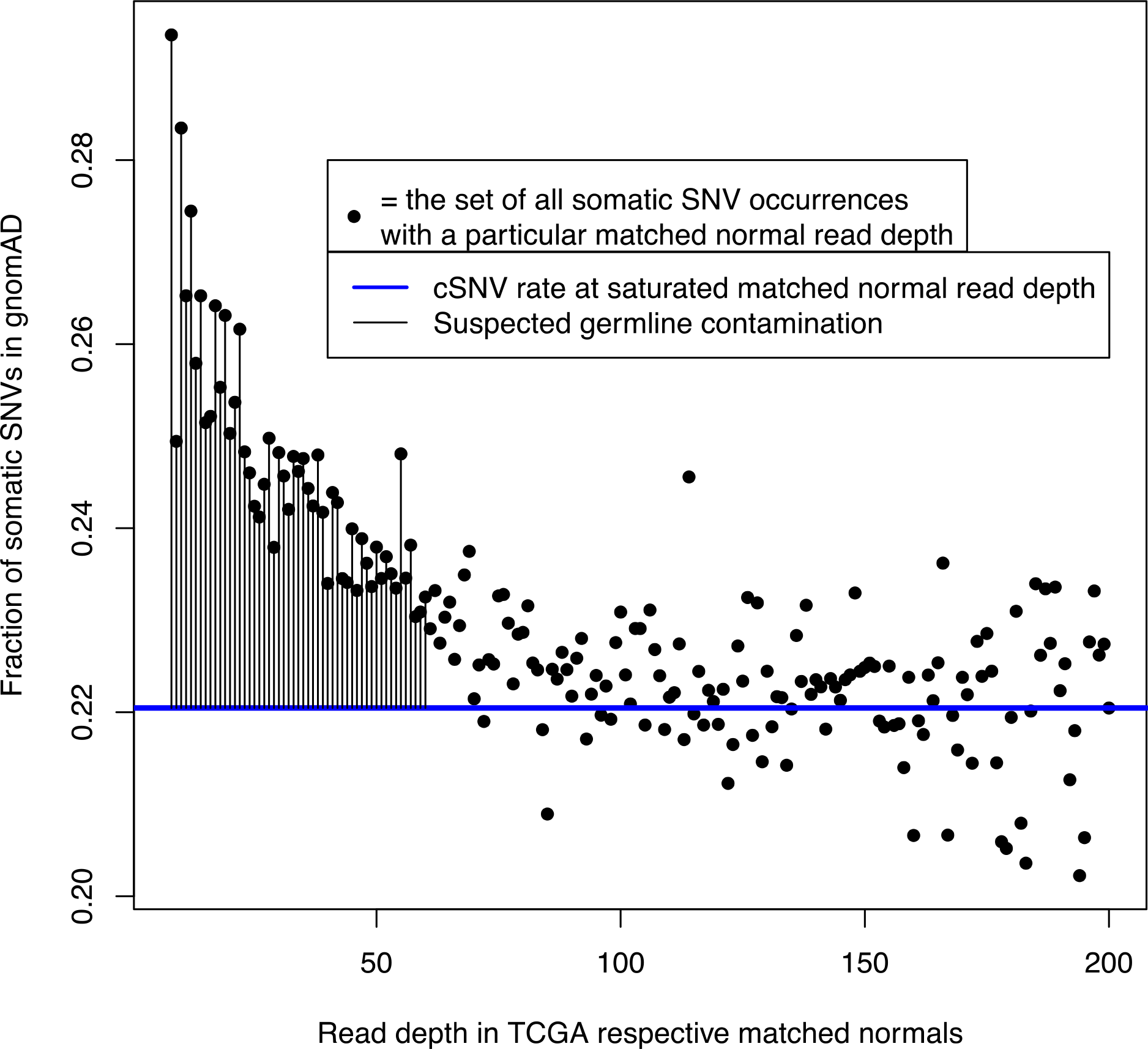
gnomAD overlap rate is highest when TCGA has a low read depth in the matched normal. Each variant instance in TCGA was called at a particular genomic site in a particular patient’s tumor sample. That patient also supplied a genomic sample from normal tissue. The read depth in the matched normal at the genomic site where a variant is called in the patient’s tumor is theoretically related to the efficiency of removal of germline variants from somatic call-sets. Along the x-axis is the matched normal read depth of a set of variants and the y-axis is the fraction of those variants that are cSNVs.

From the excess cSNV rate at TCGA sites of low read depth in the matched normal and the distribution of read depth, we estimate that there is approximately 1 leaked germline variant per strictly-filtered somatic exome, which would explain about 4% of the 268,250 observed overlapping variants. This number does not include leaked *de novo* germline events.

We next tested whether these putative germline contaminants explained some of the 5% excess of cSNVs that could not be explained by nucleotide context. We repeated the nucleotide context analysis on a TCGA subset in which all variants had 100 or more reads in the matched normal and therefore is likely free from germline contamination. If some of the 5% excess cSNVs not explained by nucleotide context are germline contaminants, then the unexplained cSNV excess should fall after this filtering step. Instead, we observed that the partition conditioned Forbes coefficient did not decrease from the value obtained on the full set (Seven Type partition conditioned score of 1.113 on high normal read depth subset, vs 1.059 on full data set). This indicates that germline contaminants are redundant with nucleotide context for explaining cSNVs. The reason for this redundancy appears to be that leaked germline variants have a more germline-like nucleotide context than true somatic variants (results not shown), such that analysis by nucleotide context already implicitly models germline contamination.

### Sequencing errors do not explain variants shared between the soma and germline

We next tested whether sequencing errors explained the cSNV rate. Some kinds of sequencing errors consistently affect the same genomic sites, such as in repetitive regions. We had removed repetitive regions from the analysis. We also excluded sites of common germline polymorphisms, which will remove any sequencing errors that consistently affect the same genomics sites. Therefore, we focused on testing for sequencing errors that arise inconsistently. TCGA includes a validation set [29] of 24,366 somatic variants from 222 uterine corpus endometrial carcinoma samples that underwent targeted resequencing and met our filters. If cSNVs result from stochastic sequencing errors, we would expect that the validation rate of cSNVs would be lower than of non-cSNV somatic variants. Instead, we find that the validation rate of cSNV and non-cSNVs of re-sequenced somatic variants are indistinguishable (99.01% vs 99.03%), indicating that false-positive stochastic sequencing errors do not explain an important fraction of cSNVs.

## Discussion

The primary goal of this study was to quantify and explain similarities in human somatic and germline mutation rates through the lens of SNV-level recurrence. We have shown that there are three times as many SNVs shared between somatic and germline call-sets than expected by independence. Given that the soma and germline involve different inheritance patterns, tissue types, exogenous exposures, chromatin states, and replication modalities, this degree of overlap was initially surprising. The substantial cross-setting recurrence suggests that there may be opportunities to usefully combine somatic and germline variant data for modeling mutation patterns, particularly in the cancer types with higher cSNV enrichments (see Supplementary Table 1).

We also show in a statistical (but not necessarily causal) sense that this excess of shared variants is mostly explained by similar mutation rates in the germline and soma of basic nucleotide contexts, especially the high rate of N[C->T]G variants. In turn, these correlated mutation rates are predicted to be a consequence of the shared exposure of the soma and germline to mutation Signature 1. Extended nucleotide context, genomic region, and germline contamination each explain a small fraction of excess shared variants.

A 5% excess recurrence could not be explained by these factors alone. Simulations propose various ways in which this could be observed; one example would be an un-modeled mutational process that increases mutation rates 4-fold on 1% of the genome. The fact that the unexplained excess of shared variants is so small relative to the explained portion indicates that the genomics community has made great strides in predicting mutation rates in both the soma and germline. Indeed, some of the excess that was unexplained by the model could presumably be explained by other discovered factors, such as strand-specificity [30], transcription factor occupancy [31], and DNA curvature [32], that were omitted from the model for simplicity. Nonetheless, the genomics community’s progress in understanding mutation rates is a matter of perspective: while we can explain recurrence at the level of individual SNVs very well, (Hodgkinson 2012)[15] found that only 40% of genomic regional differences could be explained by known factors.

Our estimate for the extent of germline contamination is higher than previous estimates (1 leaked germline event per somatic exome vs 0.1 in the literature [13]). This discrepancy could reflect differences in filtering and variant calling strategy, or it could be that our read-depth titration approach is more sensitive than other methods. Alternatively, perhaps matched normal read depth correlates with low mutation rates in the germline for reasons unrelated to germline contamination. In any case, the estimated level of germline contamination in this set of variants remains sufficiently low that it does not risk de-identifying somatic exome donors, which would require 30-80 leaked variants per sample by past estimates [33]. The larger number of variants present in whole genome samples, however, will make strict filtering more important in that setting.

The fact that *de novo* variants have a higher cSNV rate than do gnomAD variants may be an artifact to the huge sample size of gnomAD and the binarization of allele count data, which makes the average mutability of SNVs present at least once in denovo-db should be expected to be higher than the average mutability of SNVs present at least once in gnomAD. In support of this hypothesis, gnomAD variants present in 5-40 gnomAD individuals, some of which represent independent recurrences across patients, have a similarly high cSNV rate as do *de novo* variants. This artifact is a limitation in our approach, but it is necessary because we cannot distinguish independently recurring germline SNVs from germline SNVs that are identical by descent from a common ancestor (Methods).

There are various ways that undiscovered determinants of mutation rates could be important despite the small noted explanatory gap. Some determinants might have a large effect but not consistently act at the same genomic sites. Alternatively, some causal but undiscovered determinants could be highly correlated with modeled factors, making their predictive contributions redundant. Another possibility is that some important undiscovered determinants of mutation rates that increase the correlation between somatic and germline mutation rates are offset by other determinants of mutation rates that decrease this correlation. Furthermore, we may need to acquire larger data sets including of *de novo* variants before maximizing the signal from coincident SNVs, especially at non-N[C->T]G sites.

This study focused on SNVs because this is where we had the highest quantity and quality of data for recurrence analysis. Insertions, deletions, copy number changes, and structural variants may also have interesting patterns when compared between the soma and germline.

The primary focus of this study is the link between somatic and germline mutation patterns. As a whole, this analysis also relates to a more general question in genomics: in a sequencing cohort, why do we observe the variants that we do, and not others? For the detection of sites with high mutation generation rates, recurrence between human germline and human somatic samples is less confounded by common ancestry than germline-only recurrence, less confounded by positive selection than somatic-only recurrence [5], is much better-powered than recurrence analysis of *de novo* variants, and does not suffer from the alignment issues that affect inter-species recurrence analysis [34]. There are particular aspects of mutation rates that will require further investigation to explain, such as differences between the mutation rates of genomic regions in the germline genome [35]. Nonetheless, the fact that we can explain 97% of somatic-germline recurrence with known factors suggests that the genomics community has already identified many of the most important predictors of human mutation rates for single nucleotide variants.

## Conclusion

Despite their differing inheritance patterns, environmental exposures, and chromatin structure, the human soma and germline share a substantial number of variants in common. These shared variants can be largely explained by the correlated mutation rates of the basic types of nucleotide context in the germline and soma. The fact that we can explain 97% of somatic-germline recurrence with known factors suggests that the genomics community has already identified many of the most important predictors of human single nucleotide variant mutation rates more generally. This work informs and encourages the exchange of data and insights between the fields of somatic and germline genomics.

## Methods

All statistics were computed using R (version 3.5.1, R Development Core Team, 2018). All R code necessary to replicate core results will be made available shortly on the authors’ GitHub page.

### Simulations

We used simulations to benchmark the power of our SNV recurrence approach. A random subset of sites from a notional reference genome were chosen to be subject to some unknown mutational process, which either increased or decreased the mutation rate at those sites. The simulations assumed that the same sites were affected in the soma and germline and that the mutation rate multiplier of affected sites was the same in the soma and germline. Random variants were then drawn according to these relative mutation rates, and the number of observed overlaps were compared to expectations under independence. Fixed parameters were chosen to resemble the actual eligible genome size and number of somatic and germline variants; that is: 50,639,535 potential SNVs, 3,339,715 germline SNVs, and 1,309,369 somatic SNVs, implying 86,354 expected cSNVs. Two additional parameters were allowed to vary: the fraction of sites affected by the mutation process, and the impact that the mutation process had on the mutation rates of affected bases. To efficiently sweep through a broad range of possibilities, the fraction of sites affected by the mutation process took on the following values in different runs: ½^10, ½^9, ½^8, …, ½^1, 1-½^2, 1-½^3, …, 1-½^10. The mutation rate multiplier of affected sites took on the following values in different runs: 2^-7, 2^-6, 2^-5, … 2^0, 2^1, … 2^7, and all combinations of these parameters were tested. Departures from expectations were then calculated using the Forbes coefficient of association.

Notably, the simplifying assumption that the exact same sites would be affected in soma and germline to the same extent is not realistic. Nonetheless, these simulations do have real-world significance. There are conceivably many mutation processes with completely orthogonal effects on mutation rates in the soma and germline. We were not interested in modeling those processes in this study because they are not expected to lead to an excess of cSNVs and therefore do not have value for explaining cSNVs. More importantly, many real mutational processes will have correlated but not identical impacts on the soma and germline. We can conceive of these real processes as a combination of two distinct mutational processes: one with identical impacts on soma and germline and one with orthogonal impacts on soma and germline, with only the former component being relevant for explaining cSNVs and being modeled in these simulations.

### Filtering

We started with a set of all possible SNVs of the reference genome hg19 and filtered down to a conservative universe of potential SNVs. Only autosomes were included because of possible sex imbalances between data sets. To minimize artifacts due to mapping errors, we excluded sites that overlapped the EncodeDac or EncodeDuke mappability blacklists [36], that are predicted to be not uniquely alignable with 24 base pair reads, that fall in repetitive regions such as genomic super duplications, simple repeats, and microsatellites, or that are otherwise flagged by RepeatMasker [37]. To minimize artifacts due to non-uniform exome capture and coverage, we restricted sites to the Broad exome interval list, required sites to have 20 or more reads in 90% of gnomAD samples, and excluded sites in which fewer than 30% or more than 70% of the surrounding 100 bases are a G or C. To minimize artifacts related to germline contamination and sequencing error hot-spots, we removed sites with a gnomAD allele frequency of 0.1% or greater. Removing sites of common human polymorphisms also had the advantage of effectively removing discrepancies between hg19 and the human ancestral genome.

Additionally, we only included single-allelic SNVs from gnomAD graded “PASS.” For TCGA, we excluded any SNV with the filter “nonpreferredpair” or “oxog.” For denovo-db, we only included variants that were obtained through whole exome sequencing.

### Binarization

In all data sets, we mark filtered SNVs as being present or absent in a database, ignoring allele count. For gnomAD, allele count is ignored because it is not possible to robustly distinguish between recurrent germline alleles that have arisen independently in multiple gnomAD patients, vs gnomAD germline alleles that are identical by common descent from a shared ancestor. For TCGA and denovo-db, allele count is similarly binarized so as to be consistent with the gnomAD processing. This binarization leads to fewer observed excess cSNVs at the most mutable genomic sites. In one sense, this binarization is a limitation, but it has a favorable side effect of preventing outlier sites from having an outsized impact.

### Nucleotide context

We used the Rsamtools [38] and GenomicRanges [39] R packages to extract the adjacent nucleotides for each potential variant, which were concatenated around the reference and alternate allele. Reverse complements were collapsed onto a central pyrimidine; for example A[G->T]C was considered an instance of G[C->A]T since wherever one of these trimers is present on the positive strand of the genome, the other trimer occurs on the negative strand of the genome.

### Region

Genomic regions were calculated by rounding up the variant position to the nearest multiple of the bin size before concatenating with the chromosome number. For example, a variant at chr1, position 123456 would be labeled with the 100 kb bin chr1_200000.

### Germline contamination

To estimate the number of germline contaminants among cSNVs, we took the baseline cSNV rate to be the cSNV rate among somatic variants with at least 200 reads in the matched normal. Then for each matched normal read depth less than or equal to 60, we calculated the number of variants that would be expected to be cSNVs from the baseline cSNV rate. We subtracted the number of baseline-expected cSNVs from the observed number of cSNVs for these variants with low read depth to estimate the total number of extra cSNVs that may result from germline contamination.

## Authors’ contributions

WM conceived the study and performed the analysis. WM and MG wrote the paper. All authors read and approved the final manuscript.

## Acknowledgements

The authors would like to thank Drs. Monkol Lek and Joseph Chang for helpful discussions.

## Supplemental Tables

**Supplemental Table 1.** Forbes coefficients differ by cancer type

| Project | Unique Variants | Expected cSNVs | Observed cSNVs | Forbes Coefficient |
| --- | --- | --- | --- | --- |
| UVM | 968 | 64 | 371 | 5.81 |
| COAD | 107911 | 7117 | 33869 | 4.76 |
| PRAD | 16393 | 1081 | 5107 | 4.72 |
| KICH | 1475 | 97 | 479 | 4.92 |
| GBM | 33579 | 2215 | 10068 | 4.55 |
| STAD | 100215 | 6609 | 30145 | 4.56 |
| LGG | 18643 | 1230 | 5387 | 4.38 |
| MESO | 1607 | 106 | 420 | 3.96 |
| READ | 30248 | 1995 | 8782 | 4.40 |
| UCEC | 344776 | 22738 | 92393 | 4.06 |
| PAAD | 15568 | 1027 | 4226 | 4.12 |
| UCS | 4809 | 317 | 1206 | 3.80 |
| PCPG | 1297 | 86 | 309 | 3.61 |
| THYM | 1710 | 113 | 416 | 3.69 |
| ESCA | 16717 | 1102 | 3838 | 3.48 |
| CESC | 39745 | 2621 | 9255 | 3.53 |
| DLBC | 2900 | 191 | 649 | 3.39 |
| THCA | 3621 | 239 | 777 | 3.25 |
| SKCM | 198196 | 13071 | 38915 | 2.98 |
| ACC | 5118 | 338 | 999 | 2.96 |
| BRCA | 51779 | 3415 | 10105 | 2.96 |
| SARC | 12056 | 795 | 2364 | 2.97 |
| HNSC | 52421 | 3457 | 9543 | 2.76 |
| LAML | 3546 | 234 | 617 | 2.64 |
| OV | 23382 | 1542 | 3936 | 2.55 |
| TGCT | 1181 | 78 | 193 | 2.48 |
| BLCA | 71731 | 4731 | 11466 | 2.42 |
| LIHC | 23041 | 1520 | 3361 | 2.21 |
| KIRC | 11436 | 754 | 1684 | 2.23 |
| CHOL | 1236 | 82 | 171 | 2.10 |
| KIRP | 10077 | 665 | 1334 | 2.01 |
| LUSC | 91172 | 6013 | 11540 | 1.92 |
| LUAD | 97501 | 6430 | 10425 | 1.62 |

Cancer types are labeled according to the official TCGA codes [41]

